# A novel approach to single cell analysis to reveal intrinsic differences in immune marker expression in unstimulated macrophages from BALB/c and C57BL/6 mouse strains

**DOI:** 10.1101/2022.05.29.493868

**Authors:** Jeremie Breda, Arka Banerjee, Rajesh Jayachandran, Jean Pieters, Mihaela Zavolan

## Abstract

Macrophages are cells of the innate immune system that provide the first line of defense against pathogens. Their functional and morphological heterogeneity is well known, though the origin of this heterogeneity is still debated. Furthermore, while mouse strains differ in the type of immune responses that they mount to individual pathogens, the range of gene expression variation among their macrophages in the absence of a specific stimulus is not known. By applying single cell RNA sequencing we here reveal the gene expression variation in pre-stimulation macrophage populations from specific pathogen-free BALB/c and C57BL/6 mice, two mouse strains that give prototypical Th2 and Th1-biased immune responses, directed towards extracellular or intracellular pathogens, respectively. We show that intrinsic differences between the macrophages of these two strains are detectable before any specific stimulation and we place the gene expression profile of these cells within the range of variation that is measured upon *in vitro* stimulation with pro-inflammatory lipopolysaccharide (LPS) and interferon γ (IFN), or anti-inflammatory IL-4. We find that C57BL/6 mice show stronger evidence of macrophage polarization than BALB/c mice, which could explain their resistance to pathogens such as *Leishmania*. Our computational methods for analyzing single cell RNA sequencing data, controlling for common sources of stochastic variation, can be more generally adopted to uncover biological variation between cell populations.

## Introduction

The large heterogeneity in the response of macrophages to stimuli is well established [1,2]. Multiple techniques have been used to decipher the nature and cause of such heterogeneity and several hypotheses have emerged, though all invoke stochastic expression of individual genes across macrophages [3]. Some studies focused on the role of general factors such as the scaling of gene expression parameters with cell size [4], while others emphasized the role of cell surface receptors and interactions with the microenvironment in generating macrophage diversity [5]. The latter aspect is receiving renewed attention in the context of tumorigenesis, where the emergent role of macrophage subsets has started to be uncovered [6]. As in other populations of cells or organisms, the diversity of macrophages is viewed as a ‘‘bet hedging’’ strategy [7,8] for the immune system, whereby a range of cellular phenotypes is induced upon immune activation, in anticipation of phenotypic variation of the pathogen. This strategy enables the immune system to rapidly respond to diverse types of rapidly evolving pathogens [9]. An interesting system for investigating the origin and potential implications of macrophage heterogeneity consists of the laboratory mouse strains C57BL/6 and BALB/c, which naturally differ in their innate immune response. C57BL/6 mice give prototypical Th1-biased immune responses, while BALB/c mice have Th2-biased immune responses [10,11]. It has further been shown that macrophages isolated from these mice respond by producing different levels of the microbicidal nitric oxide molecule [11] and with different dynamics to stimulation [12,13].

Stimulated macrophages are grouped into two major classes, classically activated (M1) and alternatively activated (M2) macrophages, along the lines of T-cell Th1 and Th2 respectively. While the classically activated macrophage is associated with a pro-inflammatory immune response, the alternatively activated immune response is associated with an anti-inflammatory-like immune response [14]. Genes such as Nos2 and Tnfa are considered markers of M1 polarization, while Arg1 and Tgfb are considered markers of M2 polarization [15]. Increasingly many studies are shedding light on the nature of M1 and M2 polarization, and whether or not they represent two ends of a spectrum of cell states [16]. For example, previous investigations of the variability of unstimulated macrophages reported differences in the expression of M1 and M2 marker genes, even though the studies were done on the RAW264.7 cells line [2], which have been shown to have the least variation of all sources of macrophage [17]. While definitely not exhaustive, the M1-M2 paradigm provides a good framework to analyze behavior, response and activity in macrophage cells from unstimulated mice.

As methods such as immunocytochemistry or FACS are restricted to relatively few cells or cell types and to a limited number of proteins at a time [18], more comprehensive approaches, based on single cell transcriptome sequencing, could provide new insights into the intrinsic heterogeneity of macrophage populations in different mouse strains. Single cell sequencing (scRNA-seq) enables the estimation of expression levels for several thousands of genes in thousands of individual cells at a time. This, in turn, enables the identification of novel cell types, reconstruction of differentiation trajectories, and elucidation of the dynamics of responses to various perturbations [19]. Interestingly, although a number of single cell studies have been carried out in mouse model systems (e.g. [20]), a comparison of the unstimulated macrophage populations of C57BL/6 and BALB/c strains has not been done. Here we applied scRNA-seq to unstimulated peritoneal macrophages obtained from specific-pathogen-free mice, with the aim of determining whether differences are already detectable in pre-stimulation macrophage populations from these mouse strains and whether they can explain some of the response variability that is observed upon stimulation.

## Results

### Gene expression profiles identify immune cell types in the mouse peritoneum

To gain insight into the cell types that populate the peritoneum of unstimulated mice, we prepared and sequenced cells from the peritoneal lavage of 3 BALB/c mice, aiming to obtain two technical replicates from each mouse by splitting the isolated cells into two distinct pools before single cell sample preparation. One of the technical replicates for mouse #3 yielded only very few reads and was discarded. For all other replicates, we estimated the relative abundances of transcripts from all genes in every cell with a method that we have recently developed [21] and then projected these data on the first two principal components. As shown in Fig 1A, for all mice and all replicates, cells separated along the two first principal components, with one large subpopulation distributed along PC1, another one along PC2 and a smaller subpopulation of cells at intermediate PC1 coordinates. Based on the marker expression (see Table S1), the samples contained B and T cells, red blood cells (RBCs) and macrophages and thus, to focus on the macrophage population, we assigned a cell type to each cell hierarchically, starting from B and T cells, then RBCs and finally macrophages (see Methods). Determining the average expression of markers typically associated with macrophages (M), B/T cells and RBCs, as well as for other hematopoietic cell types (not shown) in each sample (individual marker distributions shown in Fig. S1), we found that indeed, B/T cell and RBC markers clearly identified specific subpopulations with high expression (Fig S2A-B). Macrophages and B cells were the most abundant cell types in the peritoneal lavage of naive mice (Fig. S2C).

**Figure 1.**
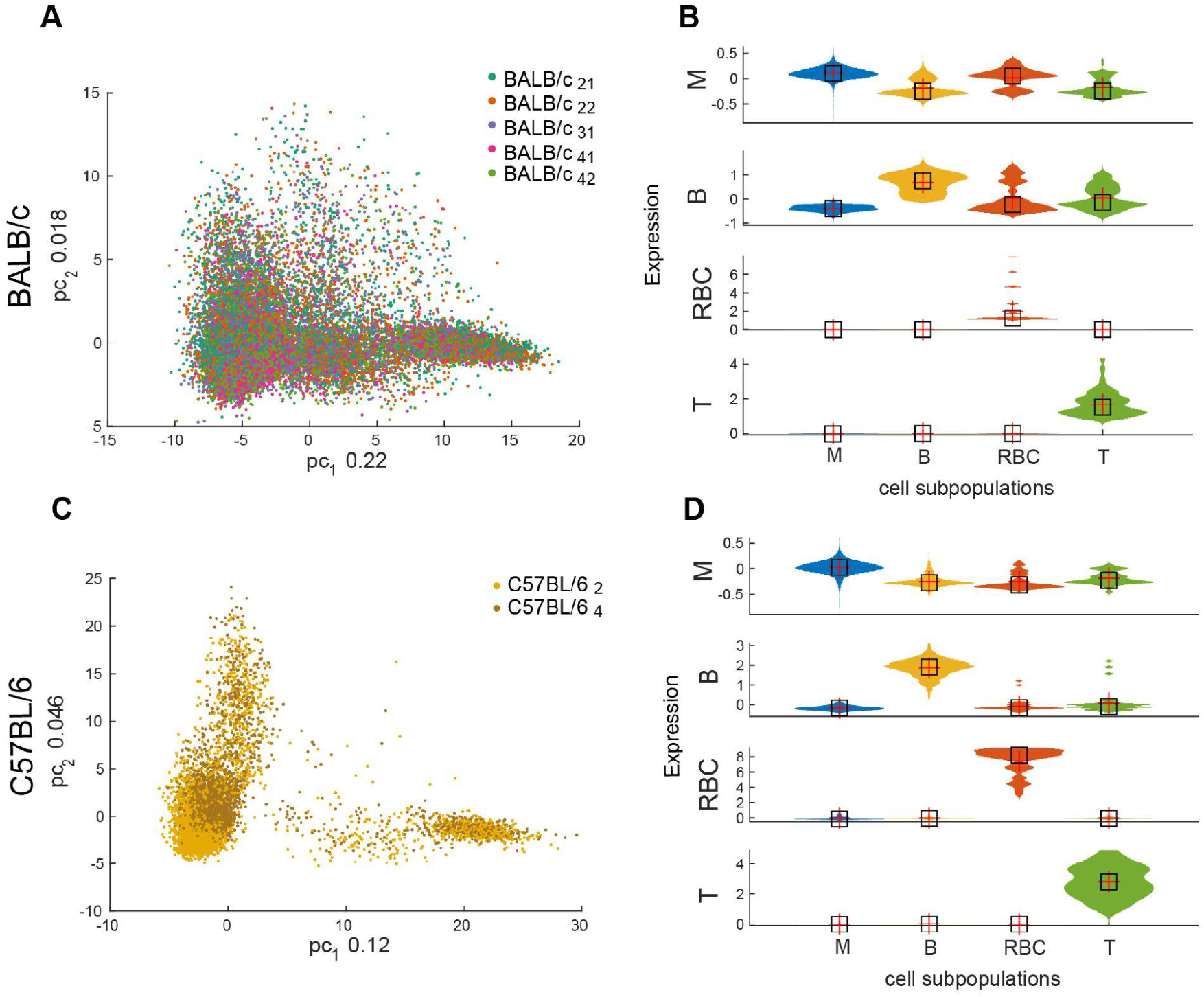
The main cell populations identified in the peritoneal lavage of mice. **A**. Projection of single cells from 3 BALB/c mice (1-2 technical replicates each) on the first two principal components of gene expression. The fraction of variance explained by each component is indicated on the axes. Individual samples are indicated by color. **B**. Violin plots showing the distribution of the average expression levels of markers of different cell types in cells from all samples from panel A. **C-D**. Similar to A-B, but for the 2 C57BL/6 mice.

We then carried out the same experiment in two distinct C57BL/6 mice. Applying the same analysis, we identified similar proportions of cells, macrophages and B cells being the two main cell types in the peritoneal lavage. The relative distribution of the cell types in the PC1-PC2 coordinate system appeared similar between the two mouse strains (Fig. 1A vs. 1C), and the distinction between the two main cell subpopulations was even clearer than for the BALB/c mice. Given the similar distributions of cells from mice of a given strain, the clearer distinction of the two main populations from the C57BL/6 mice is not due to the number of mice in the experiment but probably to the higher sequencing depth achieved for the C57BL/6 samples (Table S2). We then assessed quantitatively whether the relative proportion of cell types was similar between individual mice. Indeed, we found that within each strain, macrophages and B cells originated proportionally from each mouse, whereas the rare RBCs and T cells came primarily from individual animals (Fig. S3). Altogether, these data indicate that scRNA-seq identifies similar cell populations in the peritoneal lavage of both BALB/c and C57BL/6 mice.

### The genes that vary most across macrophages are immune response-related

We next focused on the macrophage subpopulation in the two mouse strains. To avoid ‘batch effects’ due to technical factors we centered the data from all mice of a given strain before analyzing the variability in gene expression in the dataset (Fig. 2A-B). Strikingly, in both strains, the genes contributing most to the two main directions of variation in expression across macrophages (Table S3) came from the major histocompatibility II complex or were previously associated with the M1/M2 polarization [16]. To investigate further the significance of this observation, we carried out principal component analysis using only the expression levels of these genes (Table S4). As shown in Fig. 2C-D, in both mouse strains, the expression of M1 and M2 marker defines two orthogonal axes of variation, emanating from a common center of ‘amorphous’ gene expression. Further marking the expression level of MHCII molecules in individual cells reveals a distinct subpopulation of MHCII-expressing cells, which are not specifically associated with one of the main axes of M1-M2 marker expression in Fig. 2C-D. Rather, it has been reported that the MHCII expression level in macrophages is cell cycle-dependent [22]. To validate this observation in the context of our data we determined the correlation between the average expression of cell cycle, MHCII and M1/M2 genes across cells. Indeed the average MHCII gene expression correlated better with the average expression of cell cycle genes (Pearson r = 0.26, p-value = 10^−300^ in BALB/c and Pearson r = 0.4, p-value = 10^−300^ in C57BL/6 mice) than with the average expression of M1/M2 markers (correlation between MHCII genes and M1 marker genes: Pearson r = 0.071, p-value = 1.1 10^−24^ in BALB/c and Pearson r = 0.16, p-value = 9.6 10^−61^ in C57BL/6 mice; correlation between MHCII genes and M2 marker genes: Pearson r = -0.1, p-value = 7.7 10^−49^ in BALB/c and Pearson r = 0.098, p-value = 1.2 10^−23^ in C57BL/6 mice). Thus, the expression of immune markers is highly variable in unstimulated macrophages from both C57BL/6 and BALB/c strains.

**Figure 2.**
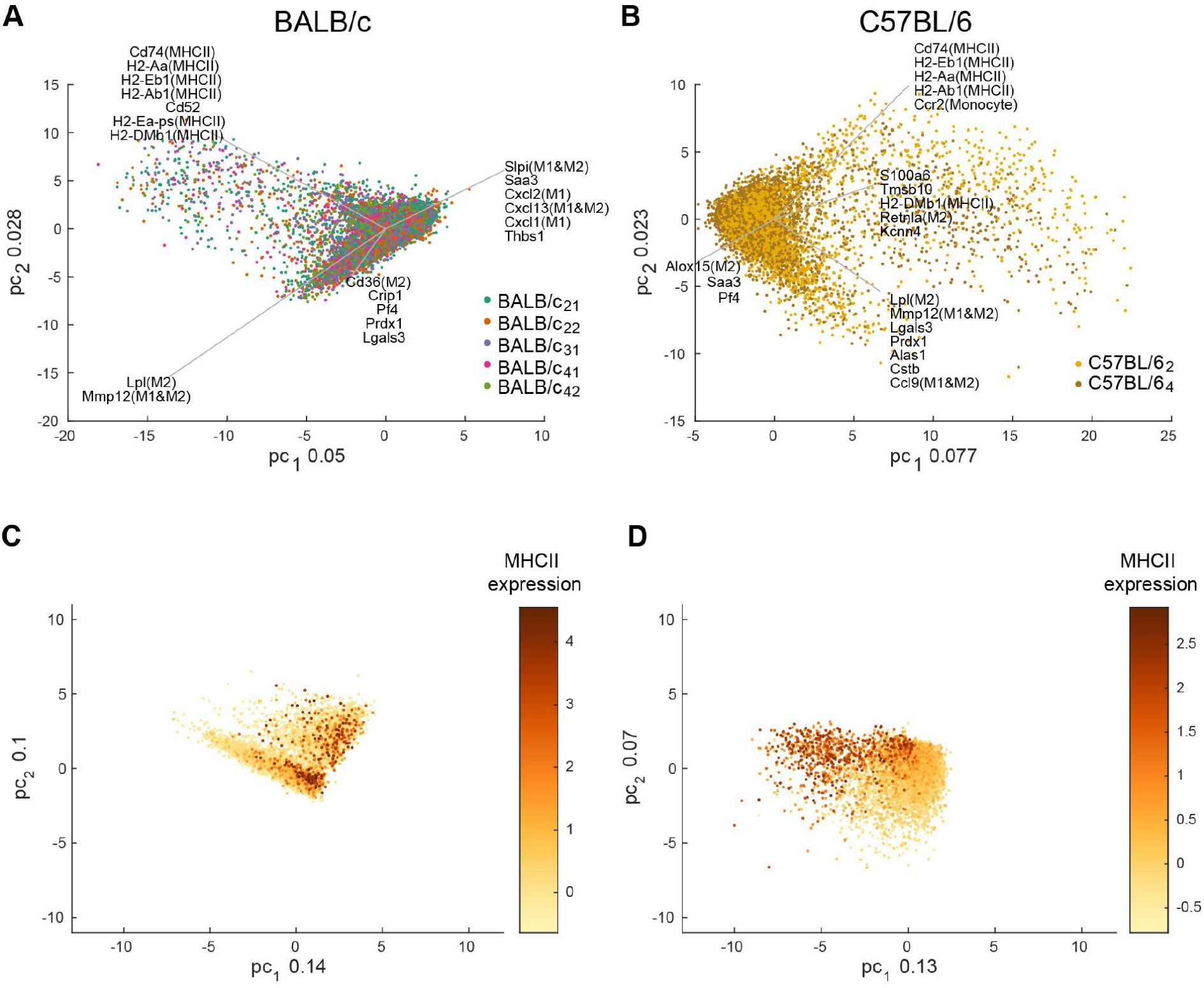
Summary of gene expression in mouse peritoneal macrophages. **A-B**. Projection of single macrophages identified in the BALB/c (A) and C57BL/6 (B) mice on the first two principal components of gene expression. The gene expression data was centered within each sample before carrying out the analysis. Individual samples are indicated by color. The genes with highest expression variability in the data set are also indicated, along with their direction of variation (see Methods). **C-D**. Similar as in A-B, but in the gene expression space of known M1/M2 markers. Cells are colored by the average expression of MHC class II genes.

### A ‘priming’ signature can be identified in macrophages from unstimulated mice

To determine whether these axes of gene expression variation are related to the M1/M2 polarization observed upon stimulation with immunological stimuli, we analyzed our data along with a previously published bulk sequencing data set of macrophages that were stimulated *in vitro* with lipopolysaccharide (LPS) and interferon γ (IFNγ), stimuli that induce M1 polarization, or IL-4, which induces M2a polarization [23]. We centered the bulk gene expression data sets on the unstimulated samples and carried out principal component analysis to identify the first two principal axes of variation in the bulk data. We then determined the distribution of each bulk sample as well as each single cell in our data in the space defined by PC1 and PC2 of the bulk data (see Methods). Bulk samples were distributed along a curved trajectory that traced well the time after stimulation. Whereas by construction most of the single cells were located at the center, along with the unstimulated bulk samples, a distinct subpopulation of cells was distributed along the axis of variation corresponding to 4-8 hours after *in vitro* stimulation with LPS+IFNγ (Fig. 3A-B). These results indicate that unstimulated macrophage populations contain cells with an expression profile that is similar to that observed after 4-8 hours of LPS+IFNγ stimulation *in vitro*.

**Figure 3.**
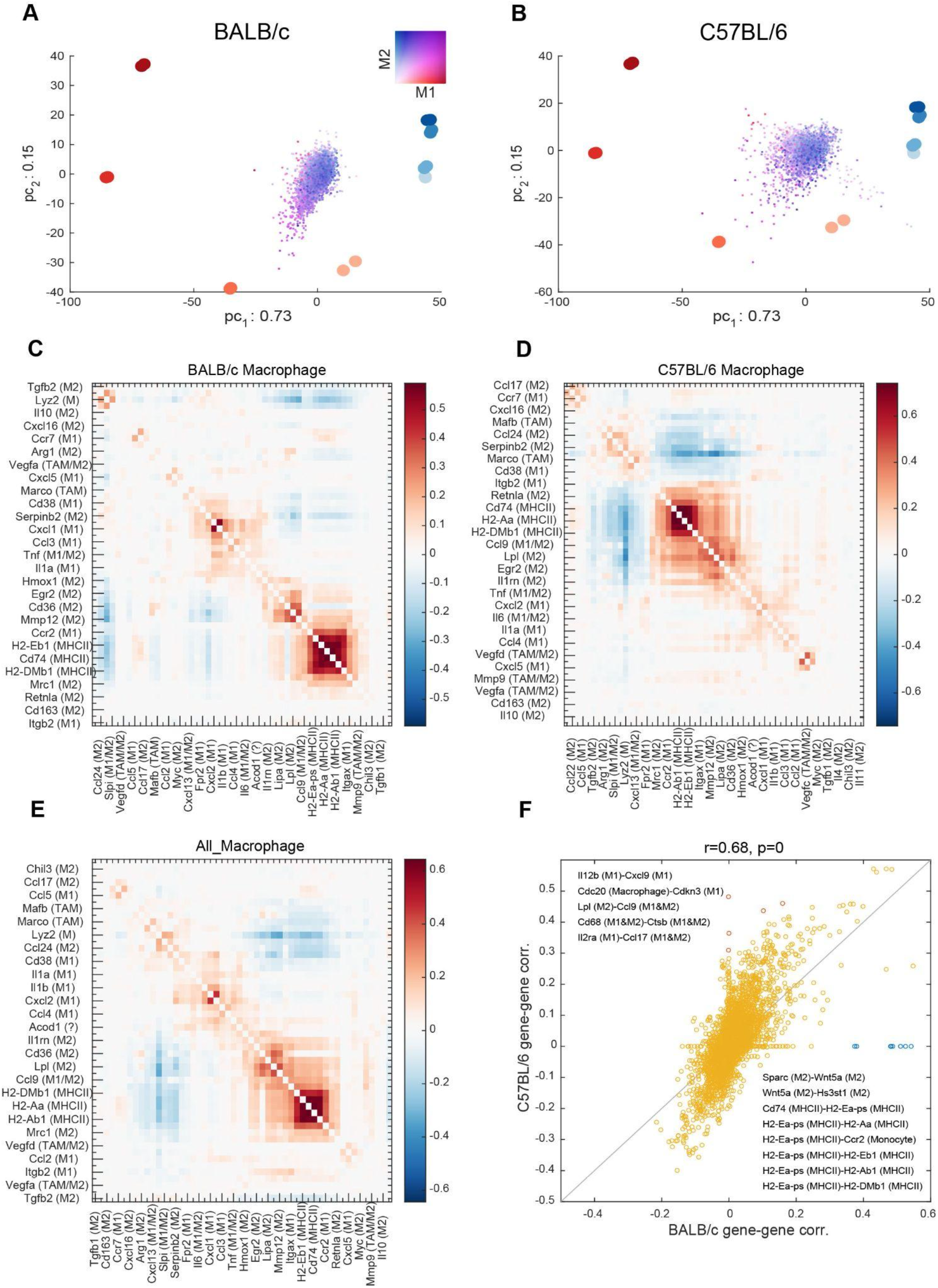
Analysis of M1/M2 bias in unstimulated mouse peritoneal macrophages. **A-B**. Projection of single macrophages from BALB/c (A) and C57BL/6 (B) mice in the gene expression space of bulk RNA seq samples from macrophages stimulated in vitro with (red), known to induce M1 bias or known to induce M2 bias (blue). For the bulk data, the shade indicates the stimulation time, 2, 4, hours. Two replicates of fold-changes with respect to time 0 (unstimulated cells) were available for each time point. The centered single cell data were anchored to the center of the bulk data, which corresponds to the gene expression state of unstimulated macrophages. The color of individual single cells indicates the relative expression of M1-M2 markers, as indicated by the 2-dimensional color legend. **C-D**. Pairwise correlation of M1/M2/MHC class II gene expression levels across individual macrophages from BALB/c (C) and C57BL/6 (D) mice. **E**. Similar, but the analysis was carried out across all cells, from both mouse strains. **F**. Scatter plot of pairwise correlation coefficients of M1/M2/MHC class II gene expression levels between the two mouse strains. Gene pairs with a relatively high difference (> 0.3) in correlation between the two strains are indicated in red (higher correlation in C57BL/6) and blue (higher correlation in BALB/c). For those gene pairs, the gene names are indicated in the corresponding quadrants.

An immediate question was whether the M1 markers are expressed in individual unstimulated cells in a coordinated manner, under the control of a common regulator, or rather independently of each other, and thus as a result of stochastic transcriptional activation of individual marker genes. To answer this question, we constructed the matrix of pairwise correlations of marker expression (Fig. 3C-E), finding that blocks of stably correlated expression emerged. The highest correlations were found among the MHC class II molecules, and also among some of the M1 markers (Cxcl1-Cxcl2, Il2ra-Ccl17), as would be expected from macrophage cells with some degree of polarization [16]. Interestingly, some M2 (e.g. Tgfbi, Lpl, Lipa, and Trem2) and M1 (e.g. Atf3, Bcl2a1d, Bcl2a1a and Psme2)-associated genes were also correlated in expression with MHCII molecules, while a set of M2 markers (Itgam, Fn1, Alox15, Ccl24 and Cd14) were anti-correlated in expression with both MHC class II molecules and the M1 markers (Fig. 3E). Interestingly, the pairwise correlations in marker expression are similar between mice (Pearson r = 0.68, p-value < 10^-300, Fig. 3F). Only a few relationships are strain-specific, indicated with distinct colors in Fig. 3F. Correlations specific for the BALB/c strain involve primarily MHCII molecules, but also Wnt5a, a ligand that promotes the containment of bacterial infections by macrophages [24]. In contrast, molecules whose expression correlates more in the C57BL/6 compared to the BALB/c mice, correspond to chemokines surface receptors and enzymes (Fig. 3F) that modulate T cell immunity (e.g. [25–27]). These data indicate that M1/M2 markers are expressed in a coordinated manner in the macrophages from both mouse strains, indicating that these correlations are induced by upstream regulators.

### Evidence of stronger polarization in unstimulated macrophages from C57BL/6 compared to BALB/c mice

We next asked whether there are genes that distinguish unstimulated macrophages from the two mouse strains, that is, whether there are genes with systematically higher expression in macrophages from one or the other mouse strain. To answer this question, we first constructed for each gene, the histogram of expression levels in macrophages from each mouse strain and the probability distribution function (Fig. 4A), taking into account the Sanity-provided error bars (from the normalization method Sanity [21] (see Methods)). This is crucial as the uncertainty of gene expression level estimates strongly depends on the expression levels themselves. Next, we constructed the reverse cumulative distributions of the gene’s expression level across cells from each mouse strain (Fig. 4B). For each expression value *x*, these distributions give the probability that a macrophage from a given strain expresses the gene of interest at a level at least as high as *x*. Traversing these two distributions from high to low expression levels *z*, we recorded the probability that macrophages from the two mouse strains express the gene of interest at a level at least as high as *z*. These points define the curve shown in Fig. 4C, with the values from C57BL/6 mice on the x-axis and those from the BALB/c mice on the y-axis. A high area under this curve (AUC) indicates that a higher proportion of cells from the BALB/c mouse showed high expression level of the gene of interest and thus the gene is more ‘specific’ for BALB/c macrophages. Conversely, low AUC indicates that higher proportions of cells from the C57BL/6 mouse showed high expression levels of the gene of interest and thus the gene is more ‘specific’ for C57Bl/6 macrophages. This analysis showed that most of the genes that are ‘specific’ to the C57BL/6 mouse are immune response genes, while the few genes specific to the BALB/c mouse encode ribosomal and mitochondrial proteins (Table S5).

**Figure 4.**
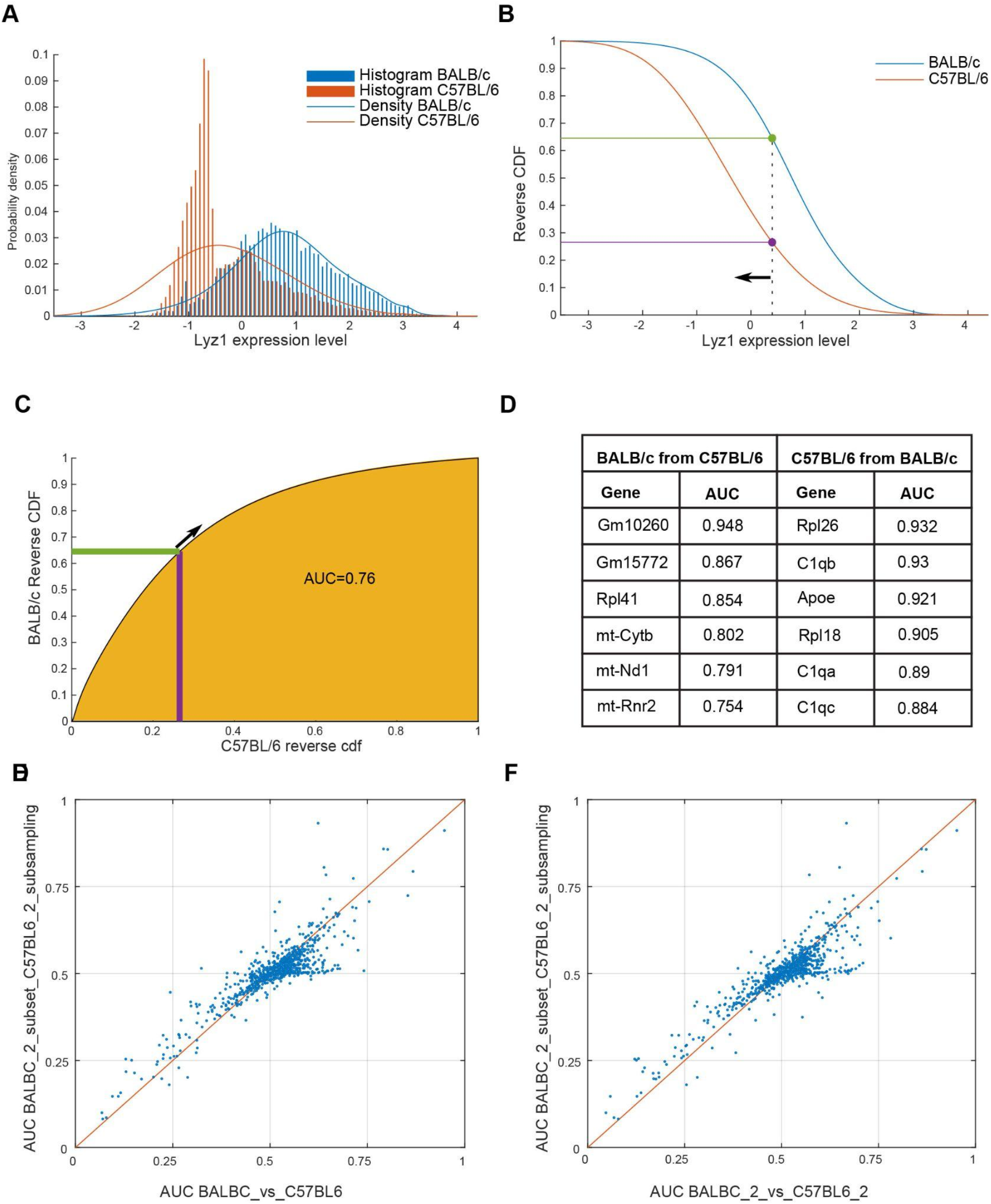
Strain-specificity of macrophage gene expression. **A**. Example inference of the gene expression level distribution across cells from individual mouse strains, taking into account the uncertainty associated with the observed read counts in single cells. The histograms show relative transcription activities [21] corresponding to inferred expression levels in individual cells, while the curves show the probability densities of relative transcription activities, calculated by taking into account the uncertainty in the point estimates of gene expression in individual cells. **B**. Reverse cumulative distributions of the same gene’s expression across cells from BALB/c (blue) and C57BL/6 (red) mice. The proportion of cells from BALB/c mice that have an expression level at least as high as the value indicated by the dashed vertical line is shown with the green and purple lines for the two strains, respectively. **C**. Inference of the AUC measure of strain-specificity of expression. The cumulative distributions shown in panel B are traversed from left to right, recording the proportion of cells from either strain whose expression level of a given gene is at least as high as indicated on the x-axis. This results in the indicated curve, the area under which is then calculated. **D**. Genes with the highest specificity of expression in BALB/c and C57BL/6 are indicated.

To ensure that these results are not due to differences in the number of cells or the number of reads per cell obtained from different strains, we carried out a randomization test. Specifically, we took the data from BALB/c mouse #2 with 7919 cells and an average of 11967 reads per cell and the data from C57BL/6 mouse #2 with 6134 cells and 37653 reads per cell and subsampled them, in order to obtain the same number of cells (6134) and a similar average number of reads per cell (11967 and 13029, for BALB/c and C57BL/6, respectively) from both mice. We then repeated the AUC analysis and the results remained unchanged (Fig. 4D-E), indicating that the specificity of gene expression that we inferred above is not due to differences in cell or mRNA capture between experiments.

The striking specificity of expression of immune genes in the C57BL/6 mice prompted us to investigate further if the degree of macrophage polarization may be higher in this strain. To answer this question, we calculated the total expression of M1 and M2 markers in each cell from each of the strains. We then traversed the expression range of M1 and M2 markers from lowest to highest, and for each expression level *x*, we calculated the ratio between the proportion of cells from each mouse with an expression level at least as high as *x* and the proportion of cells from that mouse in the entire data set. Surprisingly, we found that both C57BL/6 mice showed an enrichment of cells with high marker expression, and this was the case for both M1 and M2 markers (Fig. 5A-B). To evaluate the significance of these enrichments we randomly permuted the labels of the cells and repeated the analysis, finding that the enrichment signal disappears. Thus, these results confirm that cells from the C57BL/6 mice show higher expression of both M1/M2 markers. Then, to ensure that the enrichments are not due to differences in sequencing depth and cell numbers in the two experiments, we subsampled reads from the C57BL/6 mouse #2, to enforce an average number of reads per cell similar to those in BALB/c mice. We did not subsample the C57BL/6 mouse #4, for comparison. Repeating the subsampling 10 times, we obtained the results shown in Fig. 5C-D. Namely, the subsampling strongly reduced the enrichment of cells with high M1 marker expression, but did not affect the enrichment of cells with high M2 marker expression in the C57BL/6 mouse. These results demonstrate that the peritoneal macrophages of C57BL/6 mice exhibit evidence of polarization especially towards the M2 state, in the absence of specific stimulation.

**Figure 5.**
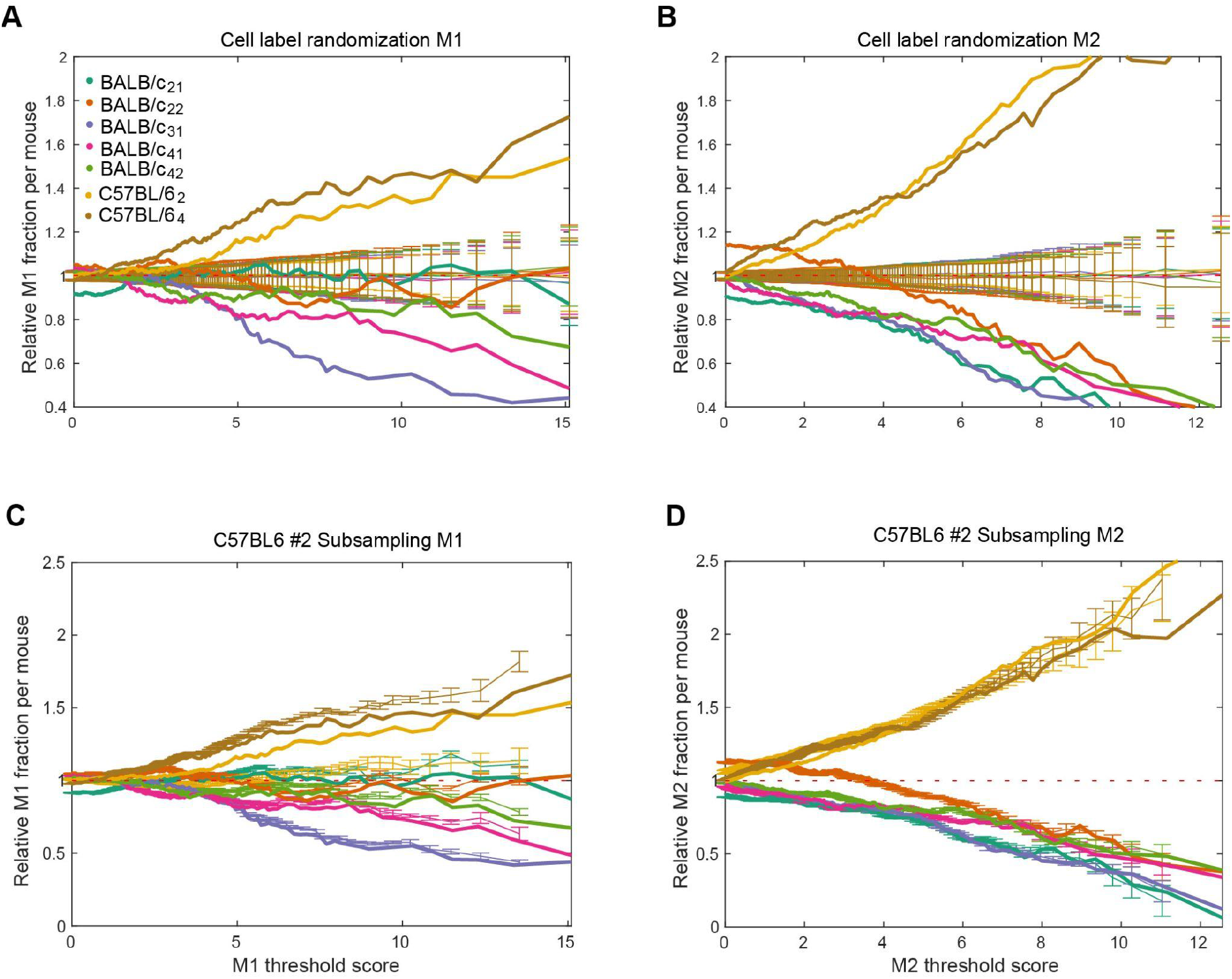
M1/M2 bias in unstimulated macrophages from BALB/c and C57BL/6 mice. **A**. M1 marker gene expression in samples from the two mouse strains. Cells from all samples were sorted by the average expression of M1 markers and then the expression range was traversed from lowest to highest, to calculate the ratio between the proportion of cells from a given sample whose expression was at least as high as the value indicated on the x-axis and the proportion of cells from that sample in the entire data set. Individual samples (thick lines) are indicated by color. For comparison, cells labels were permuted 10 times, the same analysis was repeated, and the mean and standard deviation of these results are shown in thin lines. **B**. Similar analysis for M2 marker gene expression. **C-D**. Effect of sequencing depth on the inferred degree of M1 (C) / M2 (D) bias in unstimulated macrophages. The analysis described for panel A was carried out for 10 subsamplings of the reads from the C57BL/6 mouse #2. These results are shown in thin lines with error bars, along the results on the original data, which are shown in thick lines.

## Discussion

For reasons that remain poorly understood, individuals in a population, as well as cells within an organism are not equally susceptible to infection. Macrophages are a first line of defense against various pathogens and for this reason, many studies have attempted to identify macrophage-dependent factors that can explain the variability in the response to infections. Distributed throughout the body, macrophages have different origins and functions [5]. Although simplistic, a widely-used classification has been into M1 and M2 types, also known as classically and alternatively activated macrophages [28]. Classically activated macrophages are induced by interferon gamma produced by Th1 T cells, and are involved in the response to intracellular infection such as, for example, with *Listeria monocytogenes [29]*. In contrast, alternative macrophage activation by the interleukins IL4 and IL13 produced by Th2 T cells leads to the M2 type, associated with allergies, parasitic infection and wound healing phenotype [5]. Interestingly, the laboratory mouse strains C57BL/6 and BALB/c naturally differ in their innate immune response, with C57BL/6 mice giving prototypical Th1-biased immune responses and BALB/c mice giving Th2-biased immune responses [10,11]. Consequently, these mice have different susceptibility to pathogens such as Leishmania [30].

To gain a better understanding of the intrinsic differences in the innate immune cells of these mouse strains, we have carried out single cell sequencing of peritoneal macrophages from specific pathogen-free mice. Our main results are as follows. First, by comparing single cell gene expression with the gene expression of populations of macrophages stimulated *in vitro* with either LPS+IFNγ or IL-4, stimuli known to induce M1 and M2a polarization, respectively, we found that the macrophages from specific pathogen-free animals show some degree of polarization, roughly corresponding to 4-8 hours of *in vitro* stimulation. This provides new insights into the degree of macrophage heterogeneity in mice that are not exposed to specific pathogens. Although numerous studies have been carried out on macrophages from these mouse strains, identifying raw datasets from *in vitro* stimulation experiments has been surprisingly difficult. Obtaining such data will be important for understanding the behavior of macrophages in different contexts, including the much-studied tissue-resident macrophages. Fine-grained timelines of macrophage response to various stimuli would allow improved analyses of the heterogeneity and ‘distribution of labor’ among macrophages from different sites, thus contributing to an improved understanding of immune responses and susceptibility to infections. Of great interest in this context are also technologies for single cell profiling of protein expression [31] because strain differences between the bone marrow macrophages of BALB/c and C57BL/6 mice were mainly found at the protein rather than mRNA level [32].

Furthermore, we found that the apparent polarization was due to the coherent expression of a subset, but not all, of the M1 and M2 markers. Most unexpected however was the finding that a higher proportion of cells from the C57BL/6 mouse showed increased marker expression, especially of the M2 type, which could not be explained by technical factors such as differences in the number of sampled cells or reads per cell. This suggests that the pro-inflammatory bias of C57BL/6 mice is associated with an intrinsically higher activation of immune response genes in baseline conditions. This is consistent with the observation that bone marrow macrophages of C57BL/6 mice are better equipped to deal with engulfed pathogens [32]. There are multiple cellular systems that are likely involved, including those responsible for the function of phagosomes [32], for the production and clearance of oxidant molecules [32,33], and for the direct interaction with pathogens [13]. Indeed, asking which genes can distinguish the macrophages from the two mouse strains we found that higher expression of immune response genes identifies macrophages from the C57BL/6. In contrast, we only found a few genes with higher expression macrophages from the BALB/c mouse, possibly because of the lower depth of sequencing of these samples. Analysis of single cell sequencing data poses numerous challenges [34]. Although many computational tools have been developed, methods that are anchored in the biological and biophysical nature of the data are slow to develop. However, understanding and appropriately modeling the sources of variability in gene expression levels provides increased precision in identifying genes with variable expression between cells and in the reconstruction of differentiation trajectories [21]. Our analysis took advantage of the estimates of uncertainty in gene expression levels to identify genes whose expression distinguishes macrophages from the different mouse strains. This method can be used to define markers in other contexts. This will be important for future studies because markers have generally been defined from among cell surface proteins, and the protein level gene expression correlates relatively poorly with the mRNA level gene expression, especially at the level of single cells.

In conclusion, our results provide a baseline for understanding differences in the immune response of different mouse strains, as well as a computational framework for uncovering such differences.

## Acknowledgements

This work was supported by the Swiss National Science Foundation grant #IZLIZ3_183062. The computation was carried out at sciCORE (http://scicore.unibas.ch/), the scientific computing core facility of the University of Basel.

## Methods

### Sample preparation

For extraction of mouse peritoneal macrophage, we followed a previously described protocol [35,36]. A total of 3 mice from each mouse type (BALB/c and C57BL/6) were used for the study. In short, the mice were euthanized using CO2 as per the protocol validated by the ethics committee. The abdomen of the mouse was cleaned with 70% ethanol. A small incision was made at the outer skin of the peritoneum and the inner skin lining of the peritoneum was exposed. Five ml of ice-cold 3% fetal calf serum (FCS) in 1X PBS was injected into the cavity using a 27G needle. The peritoneum was gently massaged to dislodge the immune cells in the cavity. A 25G needle was used to collect the fluid and was placed in a sterile ice-cold tube on ice. The peritoneal lavage was centrifuged at 200g for 5 min, and resuspended in RPMI-1640 medium supplemented with 10% heat inactivated FCS. The cells were plated in a 10 cm plate at 37 degrees Celsius and 5% CO2. The plates were washed 3 times after 6 hours with 1X PBS supplemented with 3% FCS to remove the floating cells. The adherent cells were scraped and used for 10X Genomics Single cell transcriptomics library preparation according to the manufacturer’s protocol.

### Reads mapping and demultiplexing

The sequencing reads (accessible from ArrayExpress, accession number E-MTAB-11814, www.ebi.ac.uk/arrayexpress/experiments/E-MTAB-11814) have been mapped and demultiplexed using *cellranger-2*.*1*.*0* (https://github.com/10XGenomics/cellranger) on the *Ensembl* genome assembly and gene annotation GRCm38, release 91 (http://ftp.ensembl.org/pub/release-91/fasta/mus_musculus/dna/Mus_musculus.GRCm38.dna.primary_assembly.fa.gz and http://ftp.ensembl.org/pub/release-91/gtf/mus_musculus/Mus_musculus.GRCm38.91.gtf.gz).

### Data normalization and analysis

The count table obtained from cellranger was normalized using the Sanity 1.0 (https://github.com/jmbreda/Sanity) [21]. The downstream analysis of the data was done using Matlab R2019b (9.7.0.1190202). The vectors of transcription activities for all genes across the cells of the different mice were initially analyzed together for all samples from a given strain. This analysis showed systematic differences in the mean activity between mice. To correct these batch effects, we centered the data within a strain, first obtaining the log2-fold differences in the transcription activity of each gene in each cell of a particular sample, and then concatenating these vectors over samples from the same strain.

### Marker-based cell type identification

The identity of each cell was assigned as follows. We explored the expression of markers for the common hematopoietic cells (erythrocytes, macrophages, granulocytes, B and T cells, NK cells, dendritic cells, endothelial and epithelial cells, platelets, hematopoietic stem cells) compiled from the literature and found expression for only B and T cells, macrophage and erythrocyte markers [37] (not shown). As many of the macrophage markers seemed to be less specific than the markers for these other cell types (Fig. S1A-B) used a hierarchical approach to assign a type to each of the cells. Specifically, we started with the B cells, for which the markers had multimodal distribution (Fig. S1B, Fig. S2A). We decomposed the average B cell marker distribution in three gaussian components, and assigned the B cell type to the cells in the gaussian component with the highest mean, using as expression threshold the value where a cell had equal probability of being part of the highest and second highest components. We then took the remaining cells and constructed a 2-dimensional scatter of T cell and erythrocyte marker expression as shown in Fig. S2B. Only very few cells had some expression of markers for these two cell types, and we chose thresholds of expression that uniquely identified cells with highest marker expression. As these cell types are extremely rare in our samples, different choices for these thresholds do not affect our results. The remaining cells were assigned the macrophage cell type.

### Identification of most variable genes

To identify the genes most responsible for the variation among macrophages we used the first two components of the gene expression data in single macrophages and multiplied the squared projection of each gene on each component by the fraction of variance captured by the corresponding component. We then summed the contributions of each gene to the two components.

### Pairwise correlations of marker genes

The correlations shown in Fig. 3C-E are Pearson correlation coefficients of the log transcription activity (see Data normalization and analysis) of each pair of macrophage subtypes marker genes (see table S4).

### Randomization of cell labels

We use the summed transcription activity of the M1/M2 markers as proxy for the respective polarization states. To assess the significance of the apparent enrichment of M1/M2-polarized cells among the cells from C57BL6 mice, we performed 100 random permutations of the cell labels. For each permutation, we computed the proportion of polarized cells coming from each mouse relative to the proportion of the different mice in the whole dataset. A cell was called M1/M2-polarized if the summed transcription activity of the respective markers was above a given threshold. We varied the threshold from 0 to the maximum summed transcription activities of markers across all cells, binning the interval of expression into 200 bins. Thus, at a threshold of 0 marker expression, the proportion of cells from a given mouse is that in the entire set of cells, and, as the threshold of maker expression increases, we observe an increase or decrease in this relative proportion depending on which strain the cells with the highest marker expression come from (see Fig 5A-B).

### Subsampling of reads and cells

As shown on Fig S4, we performed a subsampling of the reads from sample C57BL6_2 in order to match the lower reads per cell in the BALB/c samples. The average number of reads per cell being 37653 in C57BL6_2 and 12375.2 in the BALB/c samples, we randomly selected a fraction 12375.2/37568 = 0.329 of the C57BL6_2 reads directly on the fastq file. We performed 9 independent sub-samplings. To establish that the strain-specific genes that we identified (Fig. 4D) are not due to differences in the experimental procedure for sample preparation, we subsequently performed a subsampling of the BALBC_2 cells so as to match the lower total number of cells captured from the C57BL6_2 sample (see Fig. 4E-F).

### Analysis of bulk RNA-seq data from polarized macrophages

Processed data from bulk RNA-seq experiments of bone marrow macrophages stimulated *in vitro* with either LPS and IFNγ (to induce M1 polarization) or IL-4 (to induce M2a polarization) for 1, 2, 4, and 8 hours was obtained from [23], (https://www.ebi.ac.uk/biostudies/files/S-EPMC5159803/srep37446-s2.xlssheetS6.DEgene_expression). The data consisted in expression values of ∼16’532 genes whose expression differed significantly in at least one of the time points after either of the stimuli relative to that at time 0. More precisely, the table contained fold-changes of these genes at all time points relative to time 0. We centered the data for each gene and projected the samples on the first 2 principal components. We then took the single cell gene expression matrix from our experiments, centered these values across cells, and projected them on the PC1-PC2 space from the bulk sequencing, applying a scaling factor of 10, to improve visualization. We colored individual points (samples in the case of bulk sequencing and single cells in the case of our data) by the average expression of M1 and M2 marker genes (Table S4), normalized to the min-max expression interval.

## Supplementary Figures

**Figure S1.**
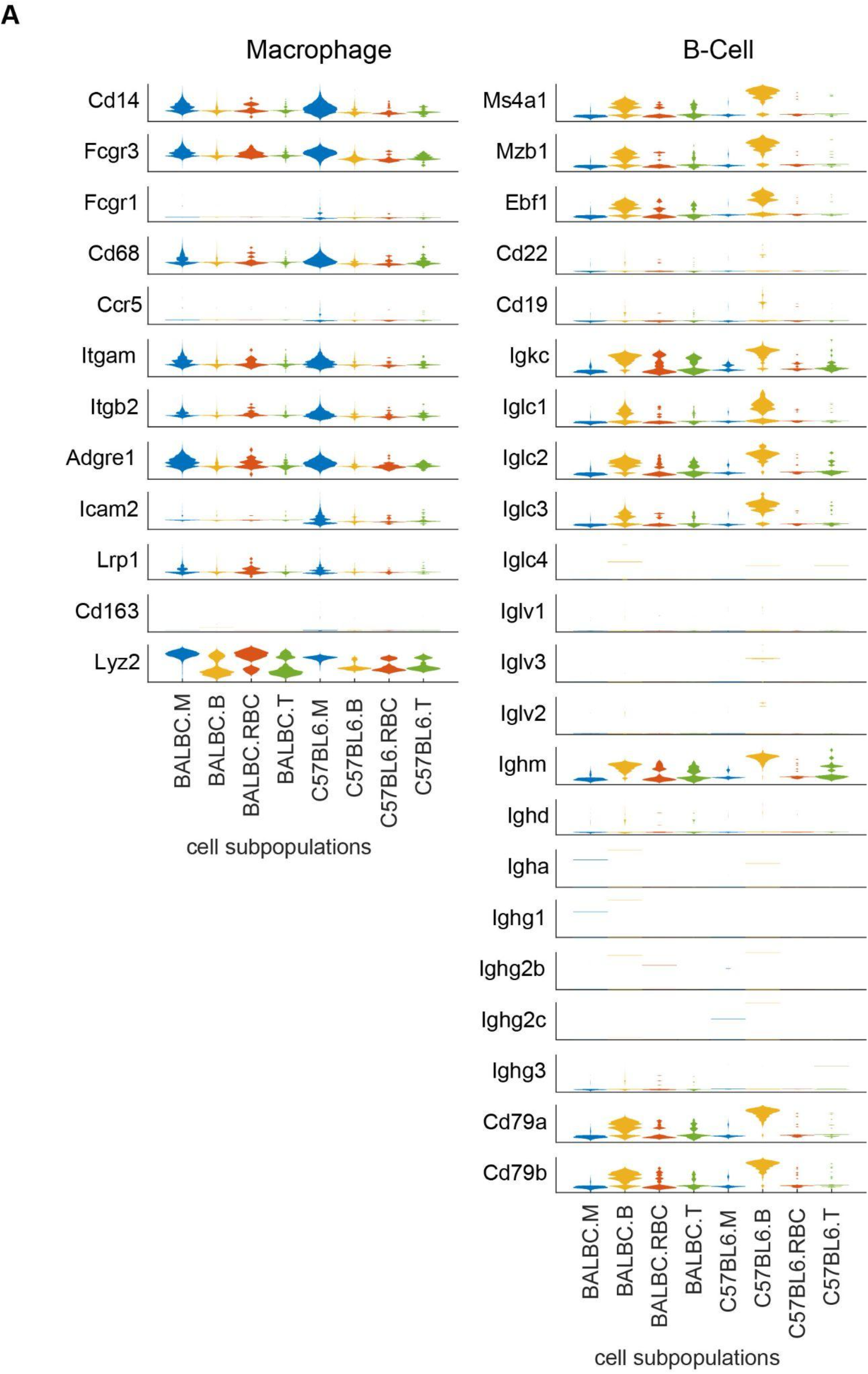
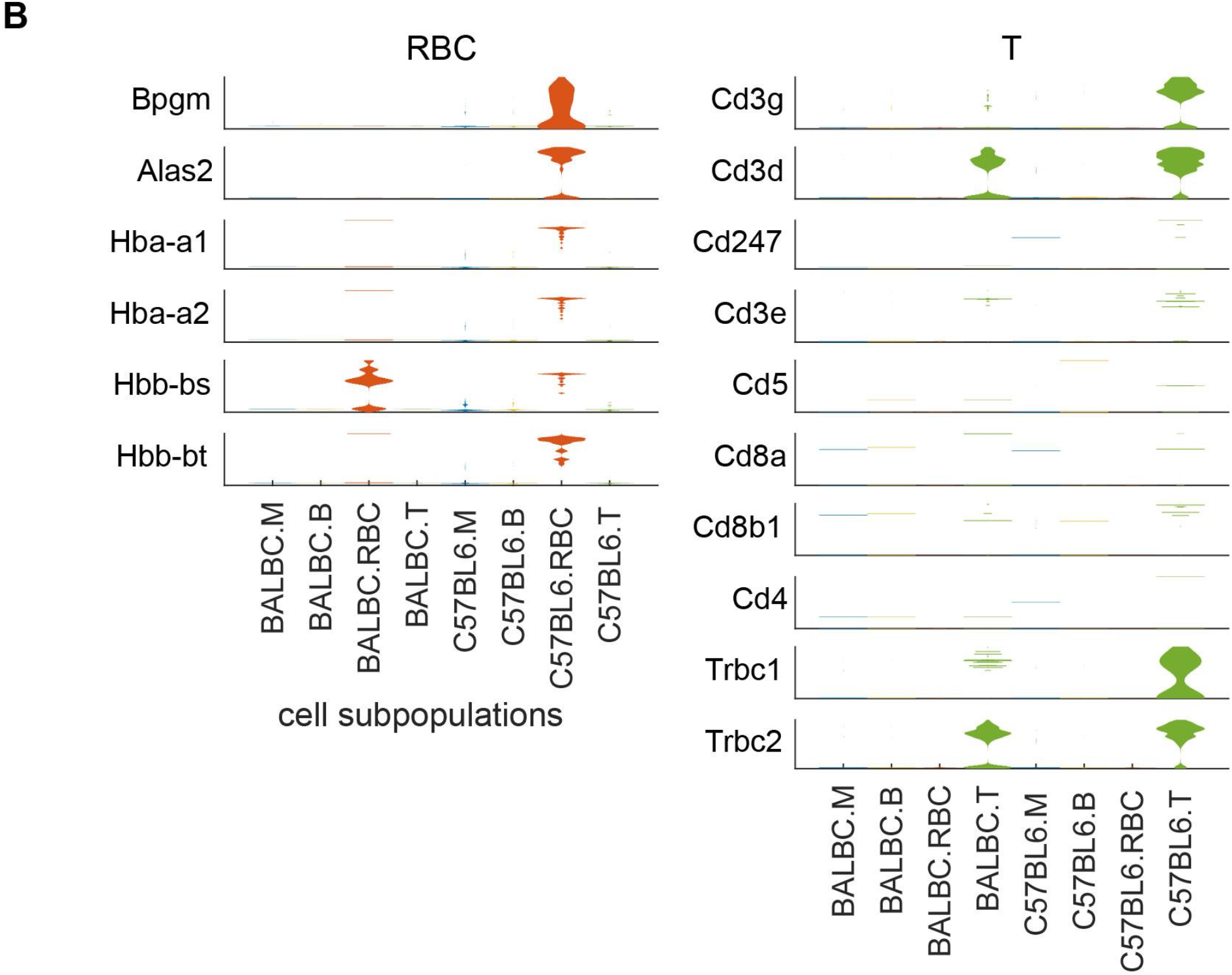
Distribution of cell type-specific makers across cells. **A-B**. Violin plots showing the expression levels of individual macrophage and B cell makers (**A**) and erythrocytes and T cell markers(**B**) in samples from all the analyzed animals. The y-axis indicates the natural log fold-change (relative to the mean across all cells) in transcriptional activity [21] for each marker (range of variation varies between markers).

**Figure S2.**
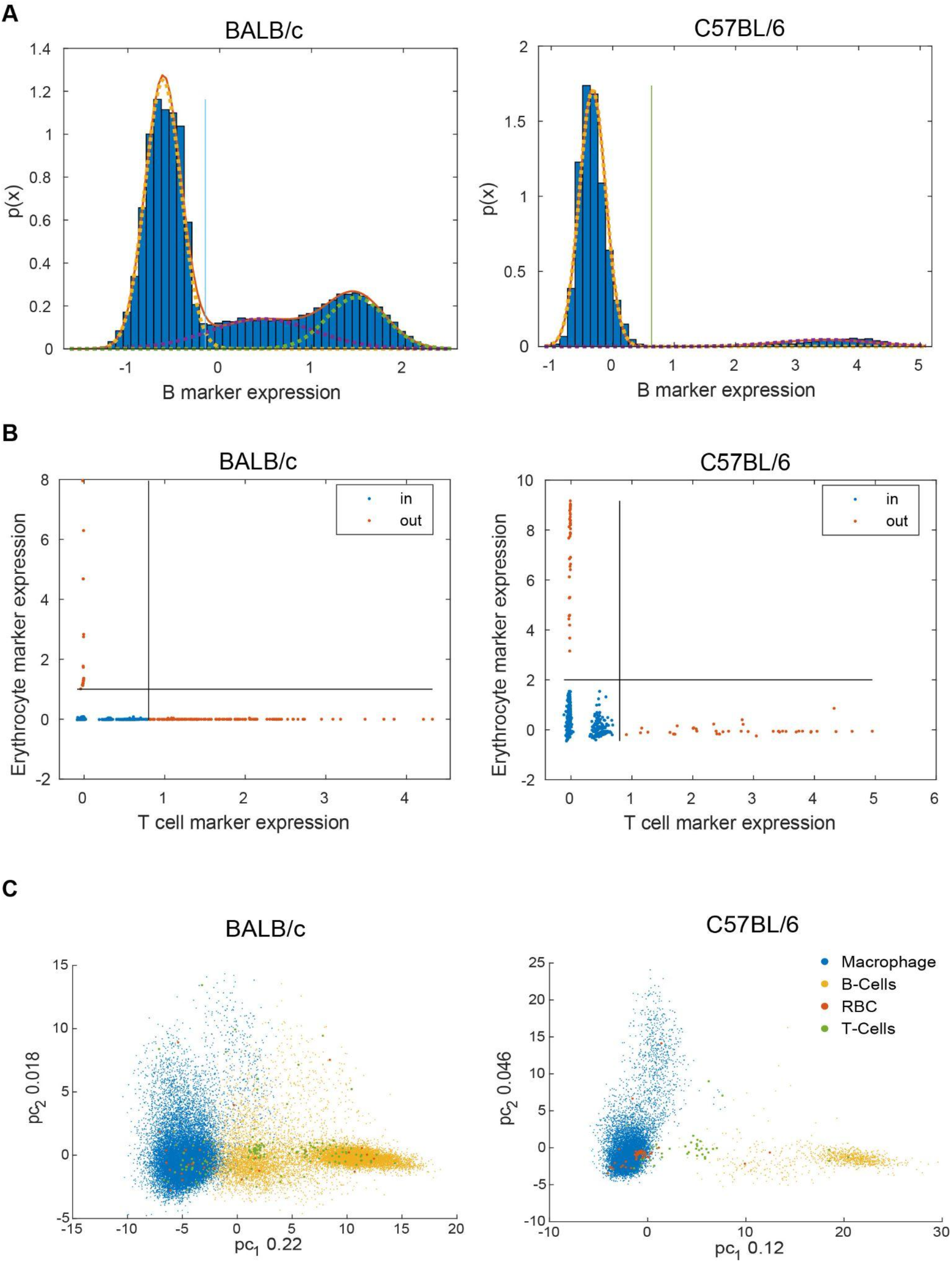
Distribution of average cell type-specific marker expression in cells from the two mouse strains. **A**. Histogram of B marker expression. **B**. Scatter plots of T cell vs. erythrocyte marker expression. **C**. Projection of all cells from each mouse strain on the first two principal components of gene expression. The cell type is indicated by the color. In each panel the left figure corresponds to BALB/c and right figure to C57BL/6 mice.

**Figure S3.**
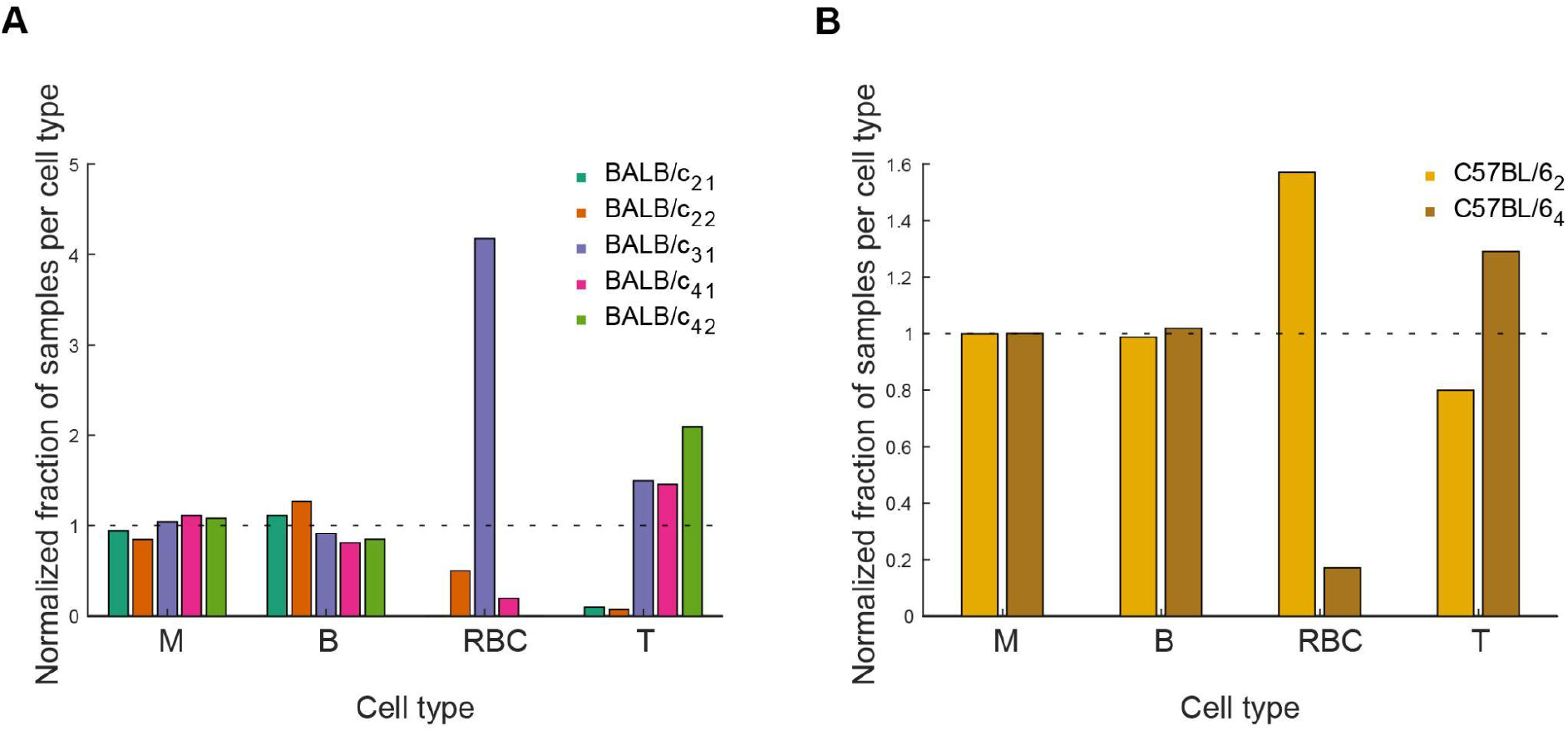
Relative proportions of individual cell types in samples from BALB/c (A) and C57BL/6. **(B) mouse samples**. The proportions of cells of a given type from individual samples were divided by the proportion of cells of any type in that sample in the data set.

## Notes

https://www.ebi.ac.uk/arrayexpress/experiments/E-MTAB-11814

### Competing Interest Statement

The authors have declared no competing interest.

